# Neuroprotective Mechanism of Ribisin A on H_2_O_2_-induced PC12 cell injury model

**DOI:** 10.1101/2023.09.27.559840

**Authors:** Xin Zhang, Mengyu Bao, Jingyi Zhang, Lihao Zhu, Di Wang, Xin Liu, Lingchuan Xu, Lijuan Luan, Yuguo Liu, Yuhong Liu

## Abstract

Ribisin A has been shown to have neurotrophic activity. The aim of this study was to evaluate the neuroprotective effect of Ribisin A on injured PC12 cells and elucidate its mechanism. In this project, PC12 cells were induced by H_2_O_2_ to establish an injury model. After treatment with Ribisin A, the neuroprotective mechanism of Ribisin A was investigated by methyl tetrazolium (MTT) assay, Enzyme-linked immunosorbent assay (ELISA), flow cytometric analysis, fluorescent probe analysis, and western blot. We found that Ribisin A decreased the rate of lactate dehydrogenase (LDH) release, increased cellular superoxide dismutase (SOD) activity, decreased the levels of tumor necrosis factor-α (TNF-α), interleukin-6 (IL-6), Ca^2+^ expression and reactive oxygen species (ROS). Moreover, Ribisin A significantly increased mitochondrial membrane potential (MMP) and inhibited apoptosis of PC12 cells. Meanwhile, Ribisin A activated the phosphorylation of ERK1/2 and its downstream molecule CREB by upregulating the expression of Trk A and Trk B, the upstream molecules of the ERK signaling pathway.

**Graphical Abstract:** 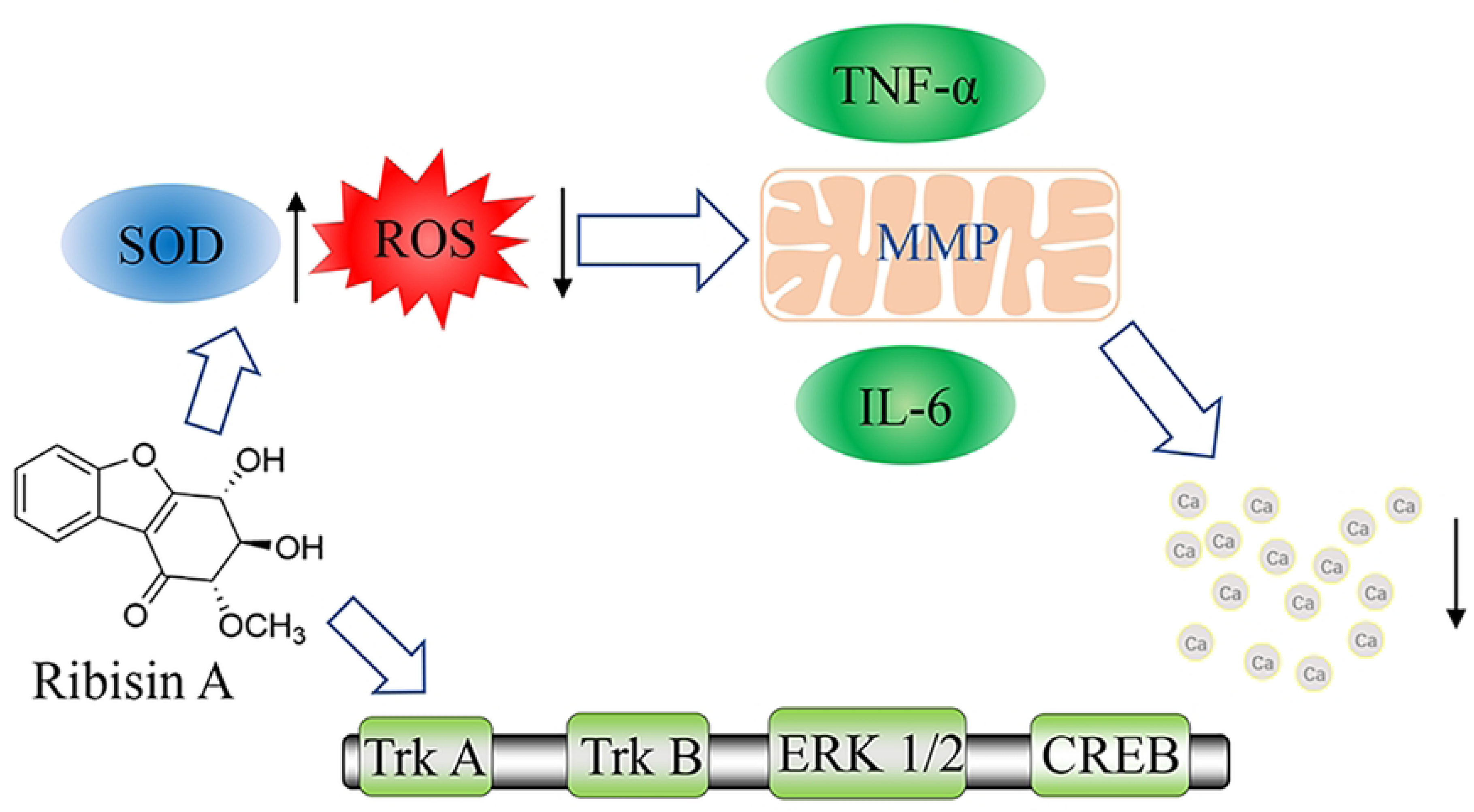

---

Alzheimer’s disease (AD) is a neurodegenerative disease with an insidious onset and gradually manifests as cognitive, language, and motor impairments, ultimately posing a serious threat to the life of the patient^1–3^. Some studies suggested that excessive deposition of A*β* is the main cause of AD pathogenesis^4,5^. They suggest that the amyloid precursor protein (APP), catalyzed by beta-site amyloid precursor protein cleaving enzyme (BACE)-1 enzyme and *γ*-secretase, eventually generates A*β*40 and A*β*42, of which A*β*42 aggregates to form oligomers that are neurotoxic and destroy healthy neurons leading to apoptosis^6,7^. In addition, there are various hypotheses on the pathogenesis of AD such as oxidative stress^8^, mitochondrial dysfunction^9^, and neuroinflammation^10,11^. However, despite recent advances in the clinical management of AD, current clinical drugs for the treatment of AD are still limited to temporarily slowing down the deterioration of symptoms and are not effective in controlling the progression of the disease. Therefore, it is necessary to develop drugs that can effectively treat AD.

There are two types of drugs commonly used clinically for the treatment of AD: one is acetylcholinesterase inhibitors (AChEI), which increase the effects of acetylcholine by reversibly inhibiting acetylcholinesterase to increase its accumulation in synapses^12^. The other is the N-methyl-D aspartate receptor (NMDAR) antagonist, which inhibits excessive activation of NMDAR to improve mitochondrial membrane potential (MMP) levels and is commonly used in the treatment of patients with moderate to severe AD^13^. However, the above two types of drugs can only treat the symptoms of AD and have no significant effect on slowing down its process. As a potential candidate pool for the development of anti-AD drugs, natural drugs have multiple advantages such as multi-target activity and low adverse effects, which are compatible with the multifactorial and diverse pathogenic mechanisms of AD^14,15^. For example, Sodium oligomannate (GV-971), a drug derived from marine brown algae, can inhibit neuroinflammation indirectly by regulating the homeostasis of intestinal flora, but is only used to delay the process in patients with mild to moderate AD^16^. Likewise, Gynostemma extract reduced the level of A*β* protein in mouse brain tissue and alleviated AD symptoms^17^. Pseudomarigold inhibited TNF-*α* production and IL-6 release, and reduced the levels of caspase 1 and 3, thereby suppressing the inflammatory response of the nervous system^18^. In addition, Chinese herbal ingredients such as Angelica polysaccharide^19^, grape flavonoids^20^, and eupulcherol A^21^ have been shown to have good anti-AD effects.

*Phellinus ribis* is a medicinal fungus of the genus Phellinus, which is widely distributed in China, Japan, Korea and other East Asian regions, and is often used as traditional medicine in China for physical weakness and senile memory loss^22^. Currently, studies on *Phellinus ribis* mainly focused on polysaccharide and benzofuranic components^23,24^. It has been reported that its polysaccharides and polysaccharide sulfation products have various biological activities such as anti-tumor, pro-angiogenic and anti-apoptotic^25–27^. Likewise, it was surprising to find significant neuroprotective effects of *Phellinus ribis*. PRG, a *β*-d-glucan isolated from *Phellinus ribis*, has shown good neurotrophic activity by inhibiting the apoptosis of PC12 cells induced by A*β*25-35^28,29^. DPRG is produced by the degradation of PRG and can play a neuroprotective role by increasing cellular mitochondrial membrane permeability (MMP) levels, decreasing cytochrome C (Cytc) protein expression, and inhibiting apoptosis^30^. Ribisin A is a new benzofuran-like compound isolated from *Phellinus ribis* that inhibits A*β*25-35-induced apoptosis in PC12 cells and significantly promotes NGF-mediated neurosynaptic growth in PC12 cells, whereas the mechanism of its neuroprotective activity is unclear^31,32^.

In this study, we constructed an *in vitro* AD model using H_2_O_2_-induced PC12 cells to investigate the neuroprotective effect and mechanism of Ribisin A. The results may lay the foundation for the research and application of new drugs for AD.

## Materials and methods

### Isolation and preparation of Ribisin A

The preparation of Ribisin A was based on previous studies^33^. *Phellinus ribis* (10.4 kg) (Licheng District, Jinan, China) was cold soaked in 5 times of methanol for 30d. The extract was concentrated to a non-alcoholic extract (200g), then dissolved in methanol and steamed dry. Silica gel column chromatography (2000.0 g, 100 ∼ 200 mesh, 5 cm × 120 cm) was then applied and sequentially eluted with CH_2_Cl_2_, CH_2_Cl_2_-EtOAc (9:1, 1:1), EtOAc, EtOAc-MeOH (9:1, 7:3, 1:1). The eluate was concentrated under reduced pressure, and the site of the benzofuran-like component was determined by NMR hydrogen and carbon (AVANCE Ⅲ-600, Bruker, German) spectroscopy. The eluate was separated and purified by Sephadex LH-20 gel column chromatography (GE Healthcare, USA), eluted with methanol, identified by TLC and then combined. The site of the benzofuran-like component was again determined by NMR hydrogen and carbon spectroscopy, and Ribisin A was separated and purified by high-pressure preparative liquid chromatography (FL-H050G, Agela Technologies, China). The column used was Innoval ODS-2 (10×250 mm, 5 um, Agela Technologies, China). Dissolve 4.96 mg of Ribisin A in 1 mL of DMSO solution and add 1 mL of sterilized ultrapure water to make 10 mmol/L of Ribisin A master batch and store at -20°C.

### Cell culture

Highly differentiated PC12 cells were purchased from Wuhan Pu-nuo-sai Life Sciences Co (Wuhan, China). PC12 cells were cultured in complete medium with 10% FBS, 1% penicillin, and 1% streptomycin at 37°C and 5% CO2, The medium was changed every other day and the cells were passaged until they grew to about 80%-90%. Cells were to be frozen and stored in liquid nitrogen.

### Choice of action mode

PC12 cells in good growth condition were inoculated into 96-well plates with 100 *μ*L per well. Then, a complete medium (RPMI 1640, Gibco, USA) containing Ribisin A and H_2_O_2_ was added to make the concentration of H_2_O_2_ per well 100 *μ*mol/L, and the final concentration of drug per well was 1, 25, and 50 *μ*mol/L, respectively, and incubated for 4h. That is, co-incubation. Pre-protection was carried out by adding 100 *μ*L of complete medium containing the above concentration of Ribisin A. After 24 h, 100 *μ*L of complete medium containing H_2_O_2_ was added and incubated for 4 h. The only difference between the restorative and pre-protective effects was the reverse order of the addition of H_2_O_2_ and Ribisin A. The concentrations of Ribisin A and H_2_O_2_ were the same in all three groups, and all were positive for Vitamin E (VE). Cell viability was detected by methyl tetrazolium (MTT) assay (Shanghai Macklin Biochemical Co., Ltd, Shanghai, China).

Cell survival %= (experimental group A570/blank control group A570)× 100%

### Detection of LDH, SOD, ROS

The treated PC12 cell cultures were collected and processed according to an lactate dehydrogenase (LDH) assay kit instructions (Applygen, Beijing, China) and the OD of LDH in the cell cultures was measured and calculated for its activity. The cell culture medium was first centrifuged and the supernatant was discarded, then the lysis buffer was added and the cells were ultrasonically crushed under the premise of ice bath. The supernatant was then separated by centrifugation at 8000 g for 10 min at 4°C, separated in a water bath at 37°C for 30 min, and the absorbance was measured at 560 nm. The superoxide dismutase (SOD) activity of the cells was calculated according to a SOD activity assay kit (Solarbio, Beijing, China). PC12 cell cultures were taken and the cells were resuspended with DCFH-DA at a final concentration of 10 *μ*mol/L. The cells were incubated for 20 min at 37°C, three times washed with serum-free medium, and intracellular reactive oxygen species (ROS) levels were measured using a flow analyzer (Beckman, CA, USA). The detection of ROS is based on the assay kit (Beyotime Biotechnology, Shanghai, China).

### Detection of inflammatory factors

The levels of TNF-*α* and IL-6 in cell culture supernatants were measured according to Enzyme-linked immunosorbent assay (ELISA) kit manufacturer’s instructions (abclonal, Wuhan, China).

### Detection of Ca^2+^ concentration

Briefly, PC12 cells were collected after treatment, washed with PBS and incubated for 20 min at 37°C under Fluo-3, AM working solution (Solarbio, Beijing, China). Then 5 times the volume of HBSS containing 1% fetal bovine serum (Sijiqing, Hangzhou, China) was added and continue to incubate for 40 min. Subsequently, cells were washed 3 times with HEPES buffer saline, and then resuspended and incubated for 10 min at 37°C. The Ca^2+^ concentration was measured by flow cytometry (Beckman, CA, USA).

### Assessment of mitochondrial mode potential

The treated cell cultures were co-incubated with the prepared JC-1 staining solution for 20 min, and then the cells were washed twice with JC-1 staining buffer. The fluorescence intensity was observed under a fluorescence microscope (zeiss, Germany), and the JC-1 monomer was detected with the excitation light set at 490 nm and the emission light set at 530 nm. The excitation light was set to 525 nm and the emission light to 590 nm for the detection of JC-1 polymers.

### Evaluation of apoptosis

The pre-treated cells were resuspended with diluted Binding Buffer, and FITC Annexin V and PI (BD, NJ, USA) were added at a ratio of 1:1 and mixed at room temperature and away from light for 15 min. 400 *μ*L of diluted Binding Buffer was added to each tube and analyzed by flow cytometry.

### Western blotting analysis

After PC12 cells were subjected to the above operation, the supernatant was discarded, washed twice with pre-chilled PBS, and the total protein was extracted by adding the prepared RIPA lysate. The protein concentration was determined by BCA protein assay (Beyotime Biotechnology, Shanghai, China). The prepared proteins were separated by SDS-PAGE gel electrophoresis (Yazyme, Shanghai, China) and transferred to activated PVDF membranes (Millipore, MA, USA). The PVDF membranes were then gently rinsed with TBST and placed in 7% skim milk powder solution and closed on a shaker at room temperature for 3 h. Afterward, the primary antibodies were added after washing with TBST and incubated overnight at 4°C. The samples were washed again 5 times with TBST, secondary antibody was added and incubated for 1 h at room temperature. Finally, the protein bands were detected using a multicolor fluorescence imaging system (GE Healthcare, USA) after infiltration with Enhanced chemiluminescence (ECL) solution (Millipore, MA, USA).

### Statistical analysis

All the experiments were carried out in triplicate at least and repeated·for three times. All the figures were expressed as the mean ± SD. Statistical analysis was performed using SPSS statistical package. Difference with P<0.05 or P<0.01 was thought to be statistically significant.

## Results

### Determination of experimental conditions

To determine the effect of Ribisin A on the proliferation of PC12 cells, we cultured cells using different concentrations of Ribisin A (1, 25, 50, 75, 100 *μ*mol/L) (Fig. 1A). The survival rates of the experimental groups relative to the control group were 105.1%, 107.4%, 105.4%, 101.2%, and 108.3% at five concentrations, respectively, indicating the proliferation effect of Ribisin A on PC12 cells in the concentration range of 1∼100 *μ*mol/L. The effect was more significant at the H_2_O_2_ concentration of 1-50 μmol/L, so this concentration range was chosen for subsequent experiments.

**Fig.1.**
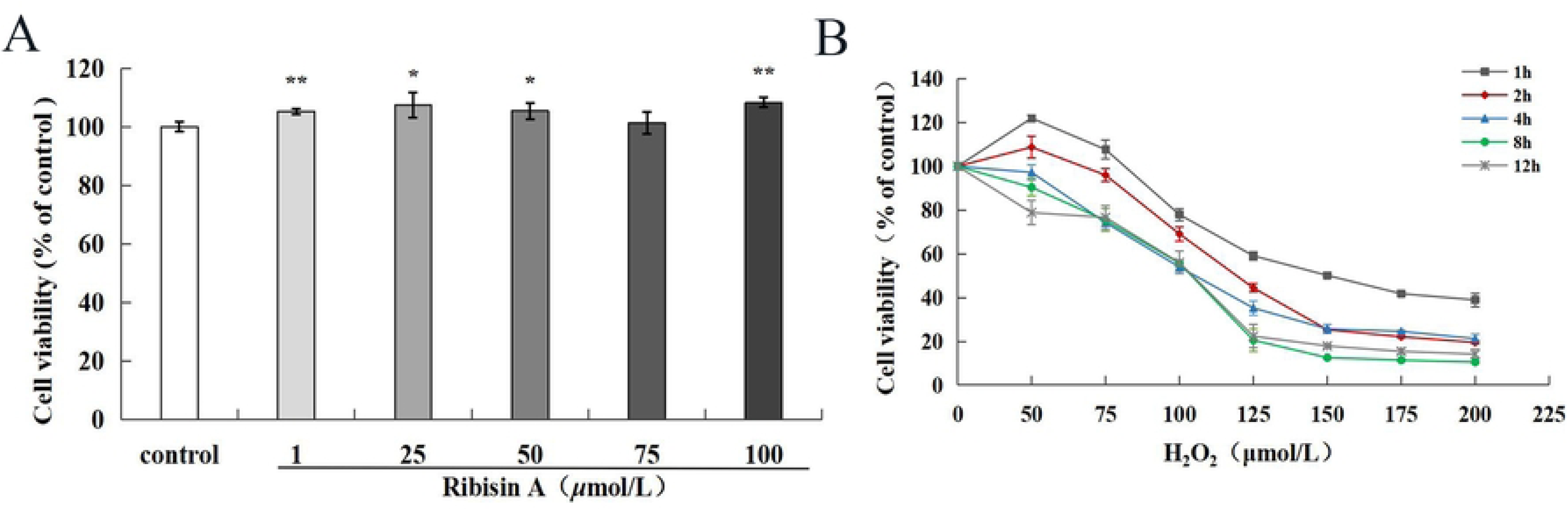
*The effect of Ribisin A on the proliferation of PC12 cells. (A) The effect of H_2_O_2_ concentration and time of action on the damage of PC12 cells. (B) The data were presented as the mean ± SD. Compared with control, * P<0.05, ** P<0.01.*

We treated PC12 cells with different concentrations of H_2_O_2_ for 1, 2, 4, 8 and 12 h, and analyzed the cell survival rate by MTT assay (Fig. 1B). We found that the number of surviving cells was 53.9% of the blank control group when 100 *μ*mol/L H_2_O_2_ was applied for 4 h, showing that the cells had been subjected to oxidative damage, but still had the possibility of rejuvenation. Therefore, this condition was considered to be the best choice for establishing a model of H_2_O_2_-induced damage in PC12 cells.

### Comparison of pre-protection, restoration and co-incubation effects

The survival rate of normal PC12 cells was significantly reduced after 4 h treatment with 100 *μ*mol/L H_2_O_2_, and the cell viability of the model group decreased significantly compared with the control group (P<0.01). As shown in the figure (Fig. 2A), PC12 cells were preincubated with different concentrations of Ribisin A before the injury, the cell survival rate was significantly higher than the model group when the concentration of Ribisin A was 25 and 50 *μ*mol/L, which could inhibit the injury of H_2_O_2_ to a certain extent.

**Fig.2.**
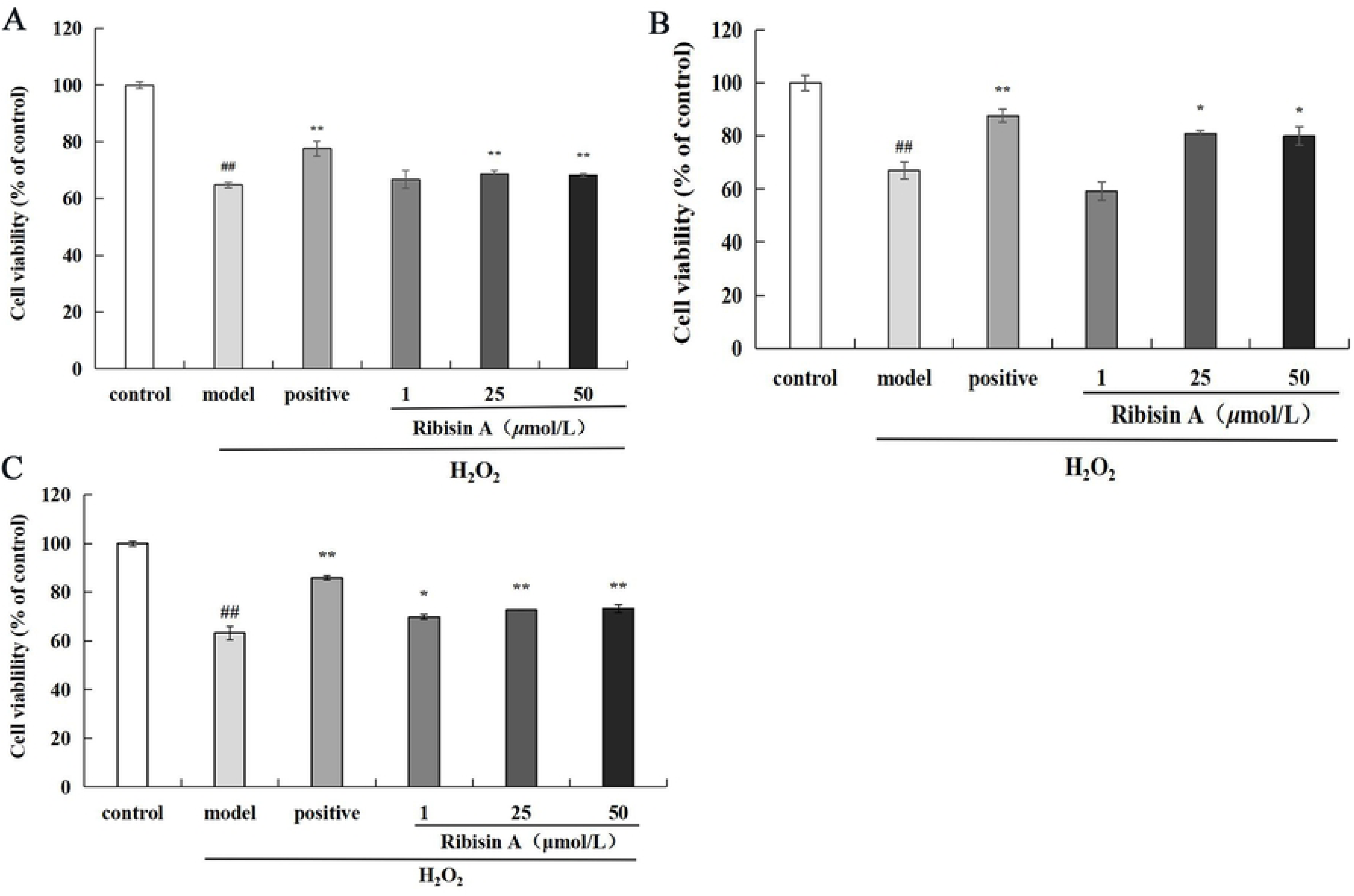
*Pre-protection of H_2_O_2_-induced damage cells by Ribisin A. (A) Restoration of H_2_O_2_-induced damage cells by Ribisin A. (B) Co-incubation of H_2_O2-induced damage cells by Ribisin A. (C) Compared with control, ## P<0.01; compared with model, * P<0.05, ** P<0.01*.

The survival rate was significantly reduced after treatment with 100 *μ*mol/L H_2_O_2_ for 4 h (P<0.01), indicating that the cells were injured by H_2_O_2_. When the concentration of Ribisin A was 1 *μ*mol/L, the cell survival rate decreased compared with the model group, but there was no significant difference (P>0.05). Whereas, the cell survival rates were 80.86% and 79.92% at concentrations of 25 and 50 *μ*mol/L of Ribisin A, respectively, which were significantly different from the model group (P<0.05) (Fig. 2B).

As shown in Figure 2C, the survival rate of PC12 cells with Ribisin A at 1, 25 and 50 *μ*mol/L was dose-dependent, with cell survival rates of 69.88%, 72.64% and 73.20%, respectively, which were significantly higher than the model group (P<0.01) (Fig. 2C). Among the three modes of action, only three concentrations of Ribisin A under co-incubation significantly increased the survival rate of PC12 cells in a dose-dependent manner. Therefore, the neuroprotective mechanism of Ribisin A in AD model will be further investigated by co-incubation in the subsequent experimental study.

### The effect of Ribisin A on LDH, SOD, and ROS levels

LDH is present in the cytoplasm and will be released into the culture medium when the cell membrane is injured, reflecting the extent of cell damage^34^. Obviously, we can find that the model group caused an increase in the LDH release rate of PC12 cells, whereas it was not significant compared to the control group according to the results. However, Ribisin A at 1, 25 and 50 *μ*mol/L reduced the level of LDH compared with the model group in a dose-dependent manner (P < 0.05) (Fig. 3A). It was shown that Ribisin A reduced the release of LDH and alleviated cellular damage.

**Fig.3.**
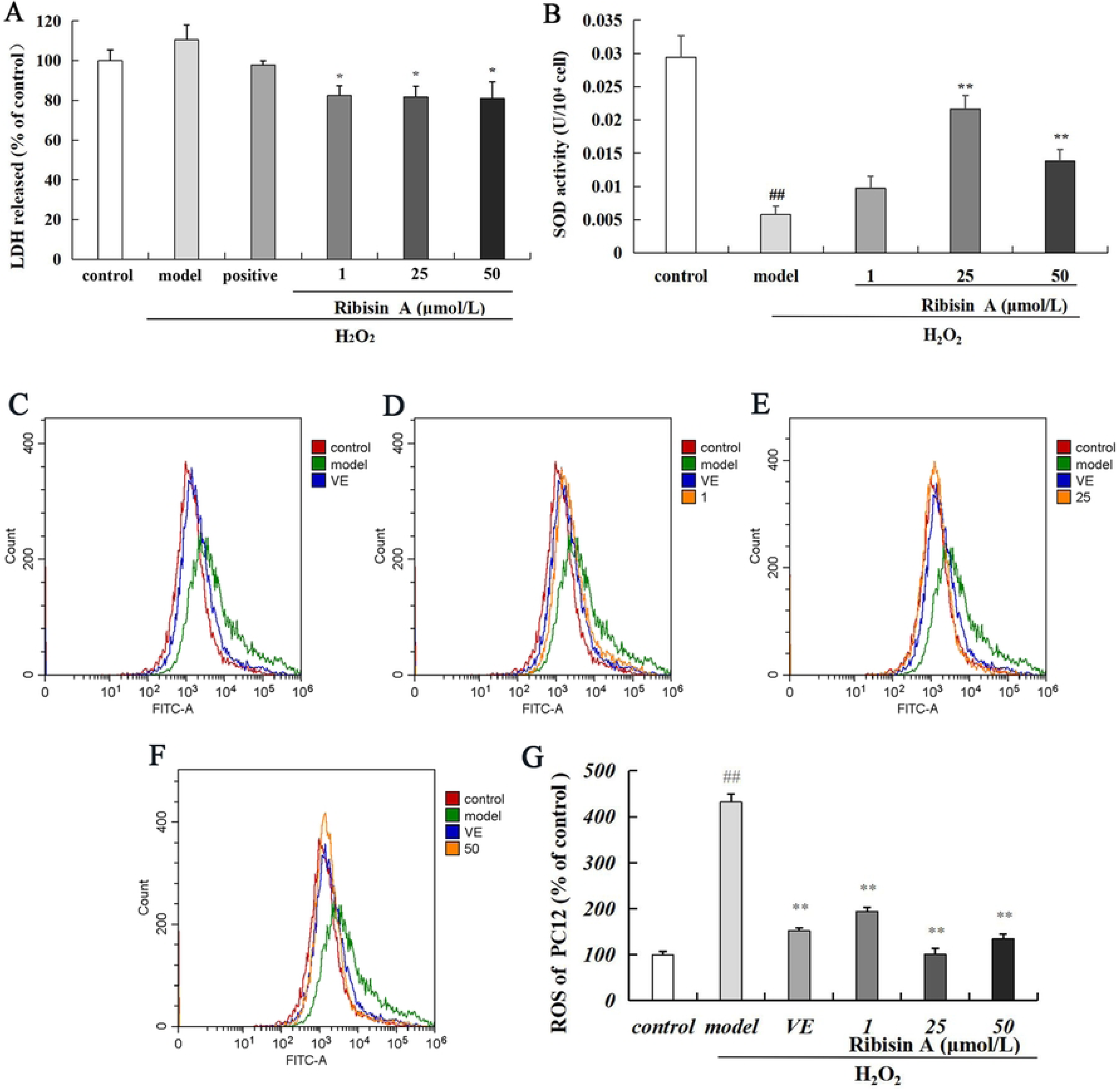
*The effect of Ribisin A on the release rate of LDH in a model of H_2_O_2_-induced injury. (A) The effect of Ribisin A on SOD levels in a model of H_2_O_2_-induced injury. (B) ROS flow histogram for control, model, and VE groups. (C) ROS flow histogram for control, model, VE, Ribisin A groups. (D-F) The relative levels of ROS in each group of cells. (G) Compared with control, ## P<0.01; compared with model, * P<0.05, ** P<0.01*.

SOD and ROS are common indicators of oxidative stress, and their levels can reflect the oxidative damage of cells^35,36^. When PC12 cells are damaged by H_2_O_2_, there is an increase in the level of free radicals due to cytosolic lipid peroxidation, and the consumption of SOD, a free radical scavenger, increases accordingly^37^. ROS is a product of normal metabolism, but cytotoxicity occurs when the level of ROS in the cell is excessively high^38^. The model group showed a significant decrease in SOD levels compared with the control group(P<0.01), suggesting an increase in intracellular SOD depletion. Compared with the model group, 1 *μ*mol/L Ribisin A increased SOD activity but was not significant (P>0.05). 25 and 50 *μ*mol/L of Ribisin A significantly increased SOD levels in the H_2_O_2_ injury model (P<0.01) (Fig. 3B). This indicates that Ribisin A can restore the activity of SOD within a certain concentration range and can counteract the oxidative damage caused by H_2_O_2_ to a certain extent. As shown in Fig. 3C, the curve of the model group was shifted to the right relative to the control group, indicating a greater intensity of cell fluorescence and higher ROS level. Compared with the model group, the curves of 1, 25 and 50 *μ*mol/L Ribisin A-treated groups were shifted to the left to varying degrees, showing a significant decrease in cellular ROS levels (P<0.01), in which 25 *μ*mol/L Ribisin A treatment showed the best results(Fig. 3D-F). The quantitative results of Fig. 3G are consistent with those shown in the flow histogram. Therefore, Ribisin A treatment can reduce the level of ROS in damaged PC12 cells, attenuating the oxidative damage caused by H_2_O_2_.

### The effect of Ribisin A on cellular inflammatory factors

Excessive accumulation of ROS can lead to the release of inflammatory factors, resulting in an inflammatory response and further toxic effects on nerve cells^39^. The level of TNF-*α* protein was significantly increased in the model group compared to the control group according to Figure 4A (p<0.01). After treatment with Ribisin A, the TNF-*α* level decreased significantly compared to the model group (P<0.01), in which 25 *μ*mol/L Ribisin A had the best effect, resulting in a decrease of TNF-*α* level to 34.38±1.25 pg/mL in the injury model. As shown in Fig. 4B, Ribisin A exerts a similar effect on IL-6 levels as TNF-*α*. Apparently, Ribisin A can protect PC12 cells by decreasing the level of inflammatory factors.

**Fig.4.**
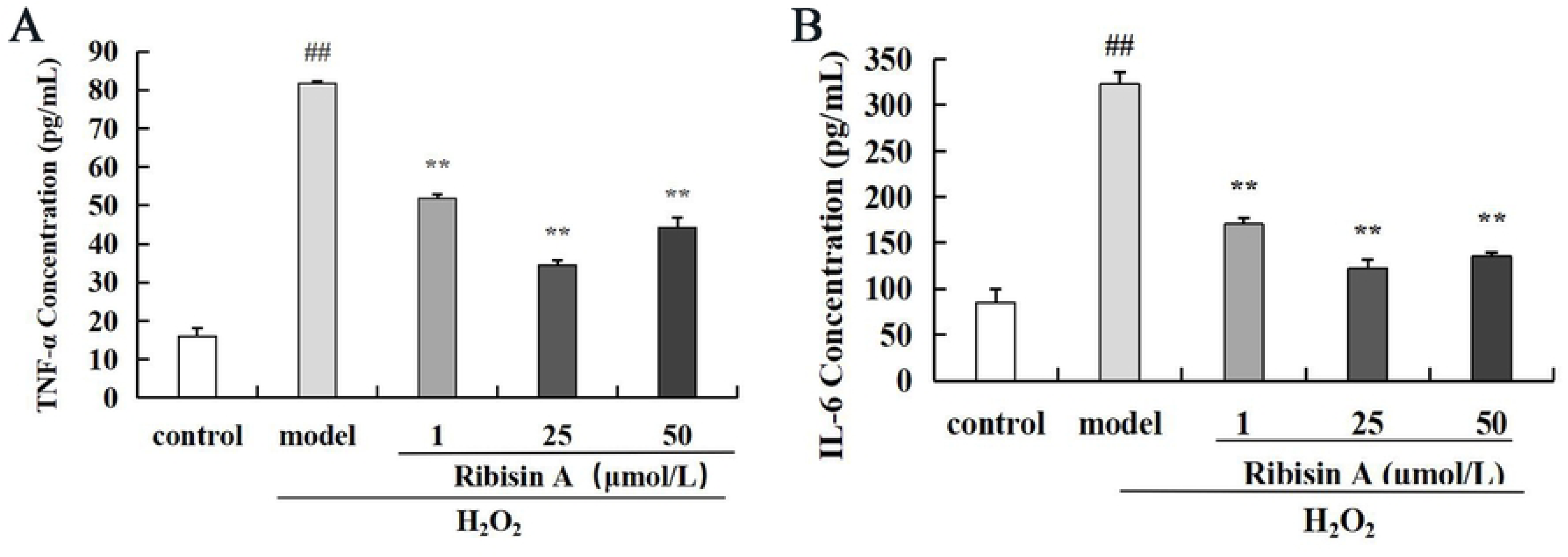
*The effect of Ribisin A on TNF-α and IL-6 Levels in a Model of H_2_O_2_-Induced Injury. (A, B) Compared with control, ## P<0.01; compared with model, ** P<0.01*.

### The effect of Ribisin A on calcium overload

Fluo-3 Am is one of the most commonly used fluorescent probes for intracellular calcium ion (Ca^2+^) concentration. It is incubated with the pentaacetoxymethyl ester of the dye and then sheared into Fluo-3 by intracellular esterases upon entry into the cell. Fluo-3 does not fluoresce in its free form, but when combined with Ca^2+^, it can produce a strong fluorescence, reflecting the level of Ca^2+^ concentration^40,41^. The histograms of Ca^2+^ fluorescence intensity are shown in Fig.5a∼e, which correspond to the quantitative results in Fig.5f. The larger the area of dark purple color in the graph, the stronger the Ca^2+^ fluorescence of the measured cells, indicating a higher level of intracellular Ca^2+^. Apparently, the fluorescence intensity was significantly enhanced in the model group compared with the control group (P<0.01), demonstrating that the Ca^2+^ concentration was significantly higher in the H_2_O_2_-induced injury model. The Ca^2+^ fluorescence intensity of the H_2_O_2_ injury model was significantly reduced by the three concentrations of Ribisin A (P<0.01). Among them, the Ca^2+^ fluorescence intensity of 25 *μ*mol/L Ribisin A group was not significant compared with 50 *μ*mol/L Ribisin A group. Overall, H_2_O_2_ stimulated the increase of Ca^2+^ concentration in PC12 cells, which could be ameliorated by Ribisin A treatment.

**Fig.5.**
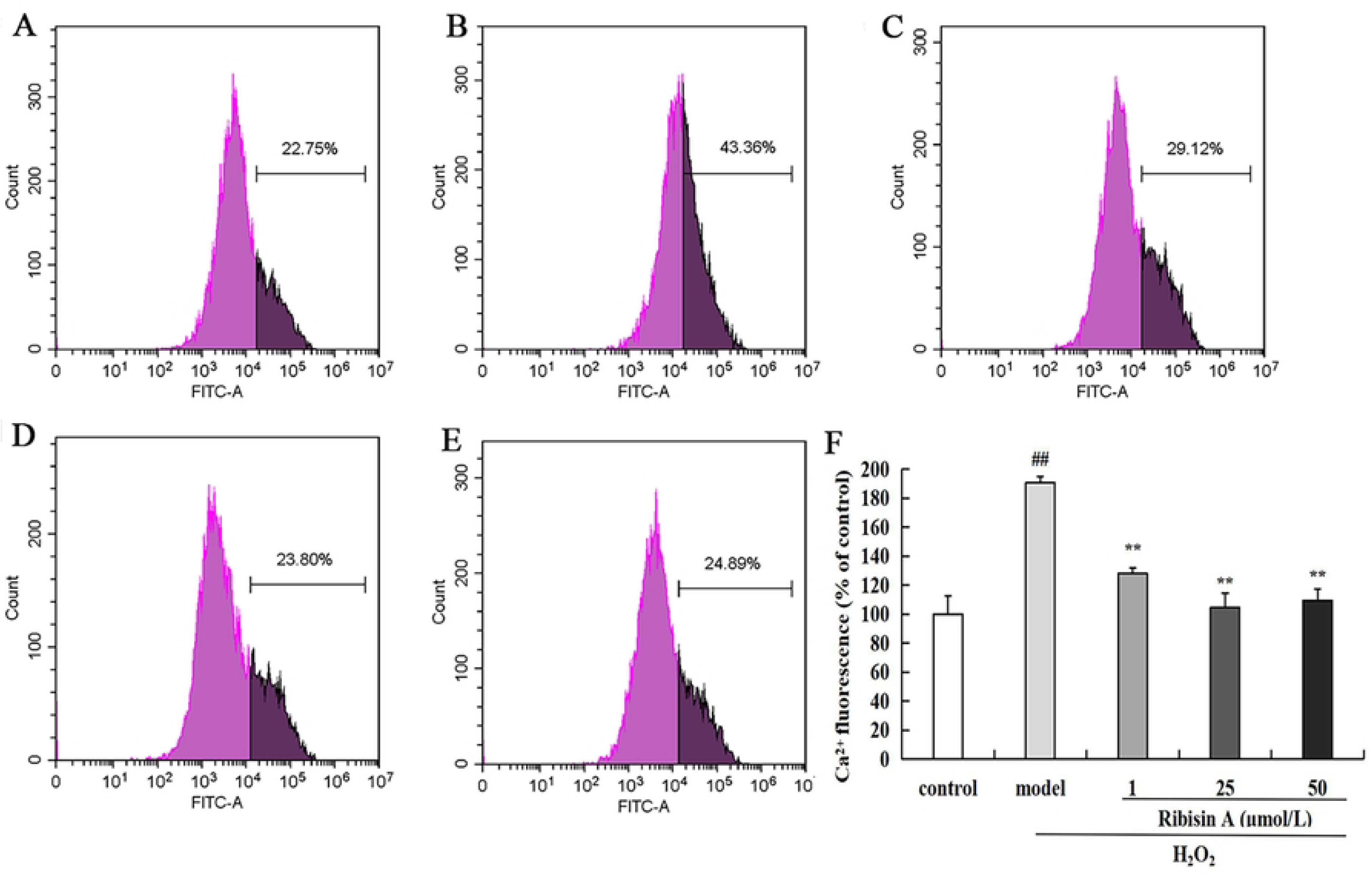
*The effect of Ribisin A on intracellular Ca^2+^ concentration. Control group. (A) Model group. (B) 1 μmol/L Ribisin A group. (C) 25 μmol/L Ribisin A group. (D) 50 μmol/L Ribisin A group. (E) Relative fluorescence intensity of Ca^2+^ in each group of cells. (F) The area of dark purple in the figure is proportional to the concentration of Ca^2+^. Compared with control, ## P<0.01; compared with model, ** P<0.01*.

### The effect of Ribisin A on MMP

JC-1 is a novel fluorescent probe whose presence status is closely related to mitochondrial function after entering the mitochondria^42^. MMP is one of the main parameter reflecting the function of mitochondria, and its alteration is an important link in causing apoptosis. JC-1 aggregates in the mitochondrial matrix to form a polymer and produces red fluorescence at high mitochondrial membrane potential. When the mitochondrial membrane potential is low, JC-1 is present as a monomer and produces green fluorescence^43,44^. According to the JC-1 fluorescence staining (Fig. 6A), the model group showed obvious green fluorescence under the fluorescence microscope compared with the control group, indicating that the MMP was low. After Ribisin A treatment, the green fluorescence was significantly weaker, suggesting an improved intracellular MMP. Quantifying the red and green fluorescence produced by JC-1 through Image J software, we found that the ratio of red/green light in the model group decreased significantly compared with the control group (P<0.01) (Fig. 6B). After Ribisin A treatment, the ratio of red/green fluorescence increased to different degrees, which is consistent with the results of fluorescence staining. It is suggested that Ribisin A can exert its neuroprotective effect by inhibiting the decrease of MMP in the H_2_O_2_-induced injury model.

**Fig.6.**
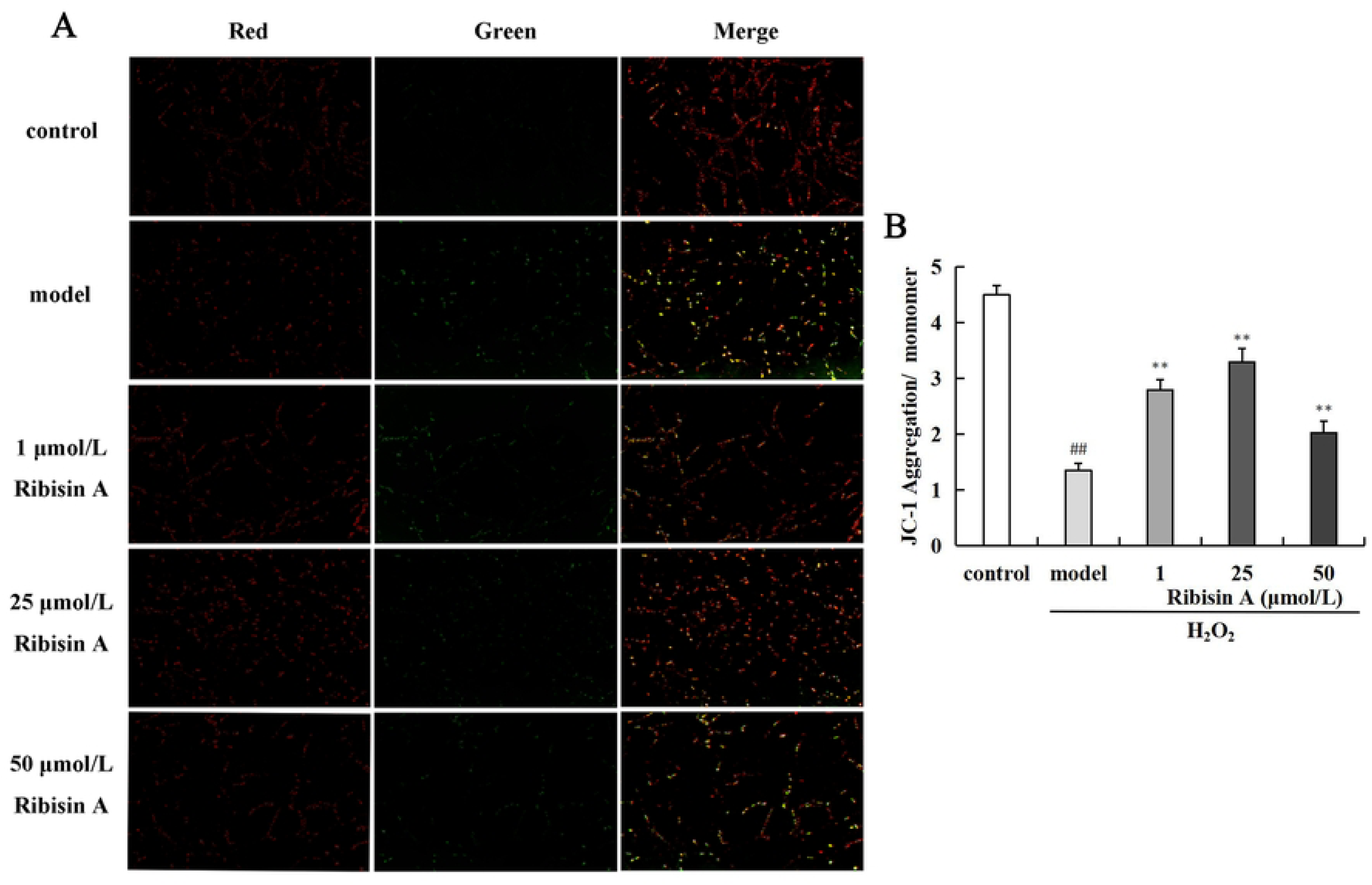
*Cell JC-1 fluorescence staining. MMP potential is high when mitochondrial function is normal, JC-1 is in polymeric state and emits red fluorescence. At depolarized membrane potential, JC-1 is in monomeric state and produces red fluorescence. (A) The effect of Ribisin A on MMP in a model of H_2_O_2_-induced injury. (B) Compared with control, ## P<0.01; compared with model, ** P<0.01*.

### The effect of Ribisin A on apoptosis

Phosphatidylserine (PS) is normally located on the inner side of the cell membrane. However, in the early stages of apoptosis, PS flips from the inner side of the cell membrane to the surface and exposed to the extracellular environment^45^. Annexin V is a phospholipid-binding protein with high affinity for PS, which binds to the cytosolic membrane through exposed PS in the early stages of apoptosis^46^. As a nucleic acid dye, propidium iodide (PI), which is normally impermeable to cell membranes, can stain the nucleus through the membranes of late-stage apoptotic cells and dead cells due to the increased permeability of cell membrane during this period^47^. So Annexin V and PI double staining can distinguish cells at different stages of apoptosis.

In the four quadrants of the flow cytometry plot, the lower left quadrant represents normal cells, the lower right quadrant is early apoptotic cells, the upper right quadrant means late apoptotic cells, and the upper left quadrant represents dead cells. As we can see in Fig. 7A-F, after Ribisin A treatment, the apoptosis rate decreased significantly (P<0.01) and in a dose-dependent manner compared with the model group, in which the treatment of 50 *μ*mol/L Ribisin A decreased the apoptosis rate to 2.82% according the quantitative results (Fig. 7G). The above results suggest that Ribisin A can exert a protective effect on PC12 cells by reducing H_2_O_2_-induced apoptosis.

**Fig.7.**
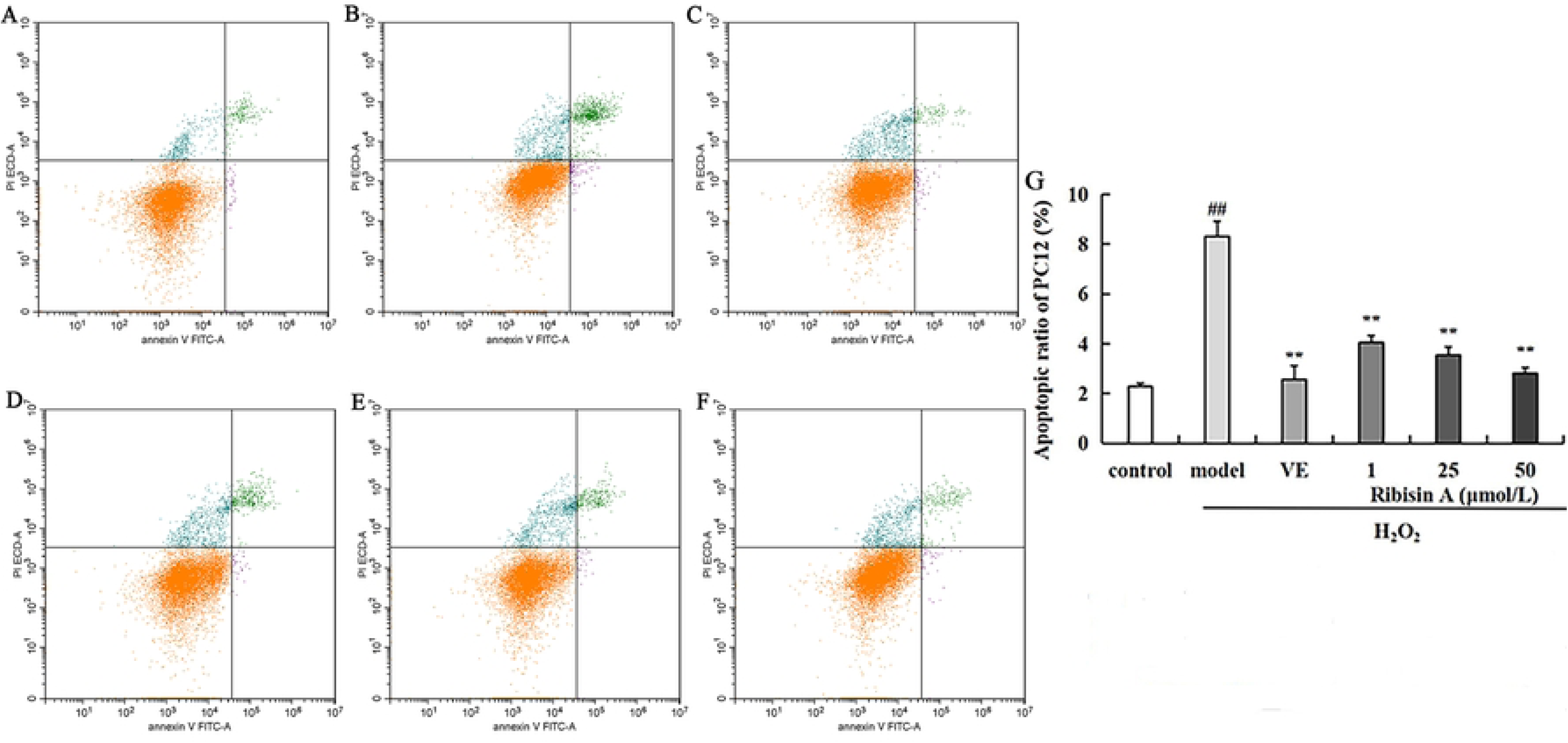
*Flow cytometry plot of the effect of Ribisin A on apoptosis in a model of H_2_O_2_-induced injury. Control group. (A) Model group. (B) VE group. (C)1 μmol/L Ribisin A group. (D) 25 μmol/L Ribisin A group. (E) 50 μmol/L Ribisin A group. (F) Quantitative results of the effect of Ribisin A on apoptosis in a model of H2O2-induced injury. (G) Compared with control, ## P<0.01; compared with model, ** P<0.01*.

### The effect of Ribisin A on the expression of ERK signaling pathway related proteins

To elucidate the molecular mechanism of the neuroprotective effect of Ribisin A, we examined the expression of ERK pathway-associated proteins by Western blotting assay and analyzed the results. As shown in Fig. 8A, the expression levels of Trk A and Trk B were reduced in the model group compared with the control group. Compared with the model group, the VE group increased the expression of Trk A but had no effect on Trk B. Meanwhile, the low, medium, and high dose groups of Ribisin A increased the expression of Trk A. The upregulation of Trk B expression was also observed at concentrations of 25 and 50 μmol/L. The quantitative results were consistent with the above description (Fig. 8B). It is noteworthy that the expression of Trk B was not increased by 1 μmol/L Ribisin A, suggesting that the upregulation of H2O2-induced decrease in Trk B protein expression by Ribisin A was effective in a certain dose range. Moreover, we found that the expression of p-ERK increased after each concentration of Ribisin A treatment compared with the model group, and the relative expression of p-ERK/ERK was consistent with the trend, in which the 25 μmol/L Ribisin A treatment group had the most obvious effect (Fig. 8C-D). These results suggest that Ribisin A can resist H2O2-induced injury by activating the phosphorylation of ERK within a certain concentration range. As shown in Fig. 8E-F, the expression of p-CREB and the relative expression of p-CREB/CREB in the model group were reduced compared with the control group, and the situation was improved to different degrees after Ribisin A low and medium dose treatment.

**Fig.8.**
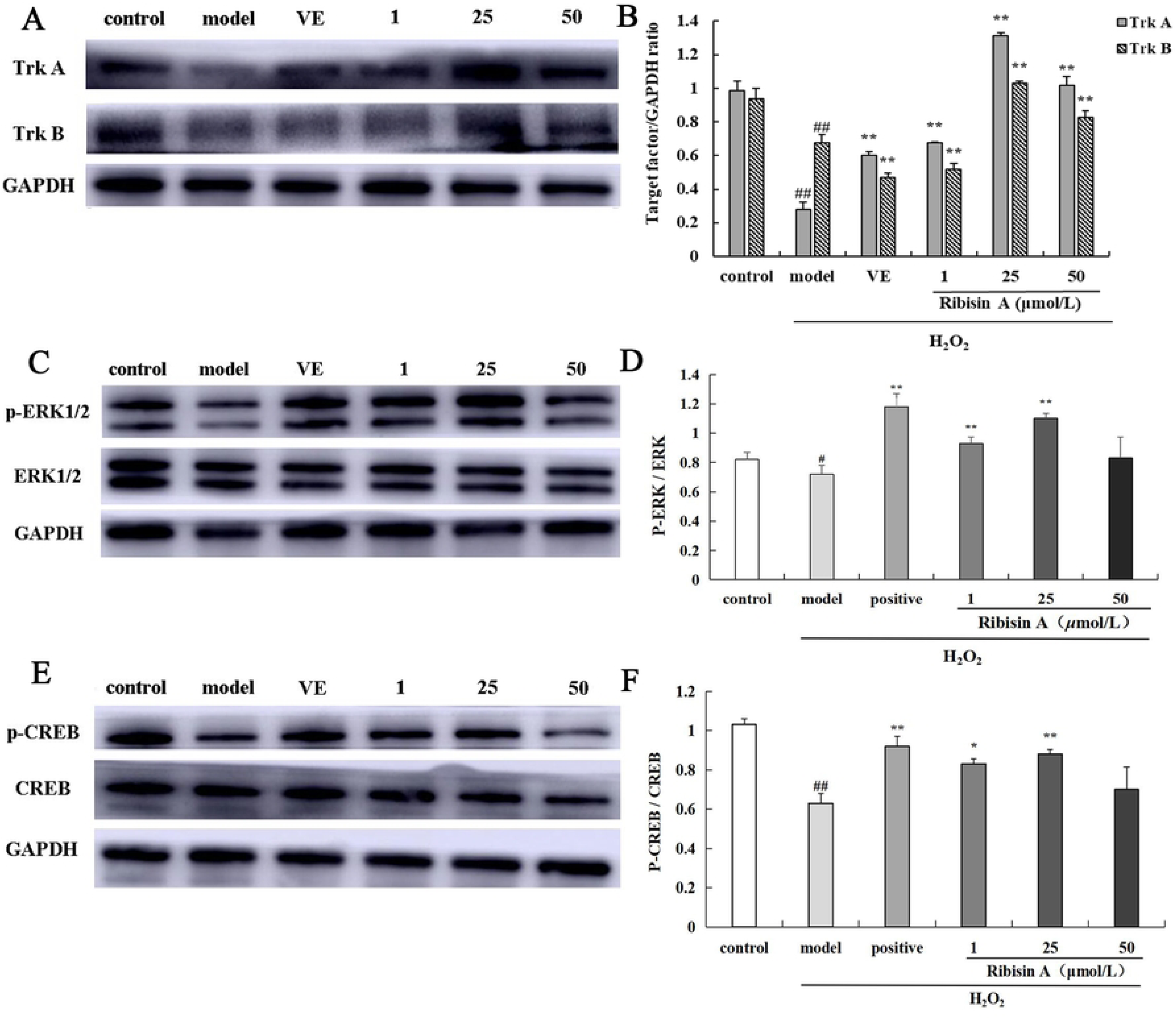
*Effect of Ribisin A on the expression of Trk A and TrkB proteins in a model of H_2_O_2_-induced injury. (A) Quantitative analysis of Trk A and TrkB protein expression. (B) Effect of Ribisin A on the expression of ERK1/2 and p-ERK1/2 proteins in a model of H2O2-induced injury. (C) Analysis of p-ERK/ERK relative expressions. (D) Effect of Ribisin A on the expression of CREB and p-CREB proteins in a model of H2O2-induced injury. (E) Analysis of p-CREB/CREB relative expressions. (F) Compared with control, # P<0.05, ## P<0.01; compared with model, * P<0.05, ** P<0.01*.

In summary, Ribisin A treatment could upregulate the expression of Trk A, Trk B, p-ERK1/2, p-CREB proteins and increase the relative expression of p-ERK/ERK and p-CREB/CREB in the H_2_O_2_-induced injury model. In this study, we showed that Ribisin A could play a protective role in H_2_O_2_-induced injury models by upregulating the expression of Trk A and Trk B proteins and thereby activating the phosphorylation of ERK1/2 and its downstream protein CREB.

## Discussion

Alzheimer’s disease is a form of dementia caused by chronic progressive central nervous system degeneration. The disease often occurs in the old age or pre-geriatric period, with dementia as the main manifestation, and the course of the disease progresses rapidly, but its exact cause is not clear^48–50^. The benzofuranic components are important active components in *Phellinus ribis* and have been shown to have a certain neurotrophic activity^51,52^. A recent study showed that ribisin C(3), ribisin G(7), and two analogues exert neuroprotective effects by targeting Keap1 and upregulating Nrf2 and its downstream target gene products heme oxygenase (HO-1) and NAD(P)H quinone reductase 1 (NQO1), activating the Keap1-Nrf2-ARE pathway^53^. Based on the above, this study investigated the neuroprotective mechanism of Ribisin A by establishing an AD model. AD models can be broadly classified into two types: cellular models and animal models. Cellular models can be subdivided into neuron-like cells and cells extracted from the brain of experimental animals directly. In this study, highly differentiated PC12 cells were used as the cell model. We found that Ribisin A can protect the AD model by reducing oxidative damage, decreasing inflammatory factor levels, restoring mitochondrial function, and reducing apoptosis by MTT assay, flow cytometric analysis, and fluorescent probes method. Western blotting showed that this protective effect may be achieved through the ERK signaling pathway.

Nerve injury caused by oxidative stress is one of the important reasons for the occurrence and development of AD^54^. As an important member of the ROS family, H_2_O_2_ has strong cell membrane permeability, which can cause cell injury or apoptosis^55^. Under the stimulation of H_2_O_2_, the excessive production of ROS causes oxidative damage to the plasma membrane, which directly causes the onset of oxidative stress^56,57^. We found that 25 *μ*mol/L Ribisin A significantly increased SOD levels, and all three concentrations of Ribisin A improved the accumulation of ROS in PC12 cells caused by H_2_O_2_ stimulation. These results suggest that Ribisin A can mitigate H_2_O_2_-induced oxidative damage to PC12 cells by increasing SOD activity and improving ROS levels within a certain range.

Neuroinflammation is an important feature of the brain in AD patients, and multiple inflammatory factors play an important role in the evolution of AD^58,59^. Excessive accumulation of ROS accelerates the release of pro-inflammatory factors, indirectly induces the production of neurotoxic mediators, aggravates the inflammatory response, and further exacerbates nerve damage^60^. In this study, we found that Ribisin A treatment could alleviate the increase of TNF-*α* and IL-6 levels induced by H_2_O_2_ stimulation, indicating that Ribisin A could exert its protective effect on PC12 cells by decreasing the level of cellular inflammatory factors. At the same time, excessive accumulation of ROS also leads to depolarization of the MMP, which induces mitochondrial dysfunction and reduces the synthesis of ATP^61,62^. As a result, Ca^2+^ in the cell membrane are limited in its transport function due to insufficient energy, resulting in calcium overload^63^. In addition, the mitochondrial apoptotic signaling pathway will be activated, leading to cell apoptosis^64^. This part of the experiment observed that cell apoptosis, reduction of MMP, and Ca^2+^ overload were observed in H_2_O_2_-induced PC12 cells. Ribisin A could counteract the injury of H_2_O_2_ on PC12 cells by significantly ameliorating these phenomena.

Previous studies have shown that different drugs can achieve neuroprotective effects through different signaling pathways, and the angles to achieve this effect are diverse. Tetrahydrocurcumin inhibits cell cycle arrest and microglia apoptosis via the Ras/ERK signaling pathway^65^. Prodigiosin attenuates oxidative injury, neuroinflammation, and apoptosis in hippocampal tissue through the Nrf2/HO-1/NF-κB signaling pathway^66^. Lactobacillus plantarum DP189 exerts neuroprotective effects by inhibiting tau protein hyperphosphorylation via PI3K/Akt/GSK-3*β*^67^. The above mentioned effects are all important ways to achieve neuroprotective effects. Besides, ERK signaling pathway is also closely associated with neuroprotection. ERK is closely linked to learning and memory functions and is essential for signaling from cell surface receptors to the nucleus^68^. ERK is normally localized in the cytoplasm, but after ROS stimulation, the phosphorylated product p-ERK is generated and translocated from the cytoplasm to the nucleus. It has been demonstrated that nerve cell activity decreases with the inhibition of the ERK pathway and neurological impairment will occur accordingly, suggesting a possible association between the ERK signaling pathway and neuroprotection^69–71^. The results of this study showed that the expression of p-ERK in the model group was inhibited, p-ERK/ERK was significantly reduced, and the expression of upstream ERK proteins Trk A and Trk B and downstream proteins p-CREB were also significantly decreased compared with the untreated group. The relative expression of p-ERK1/2 protein and p-ERK/ERK were significantly increased after 25 *μ*mol/L Ribisin A intervention, and the expression of Trk A, Trk B and p-CREB protein were increased. From the above results, we can tentatively conclude that the molecular mechanism of the protective effect of Ribisin A on the H_2_O_2_-induced injury model may be achieved by upregulating the expression of Trk A or Trk B, the upstream protein of the ERK signaling pathway, which in turn activates the phosphorylation of ERK1/2 and its downstream molecule CREB. (Fig.9) Therefore Ribisin A has unlimited potential in the treatment of AD, but future animal models are needed to further discuss its neuroprotective mechanisms.

**Fig.9.**
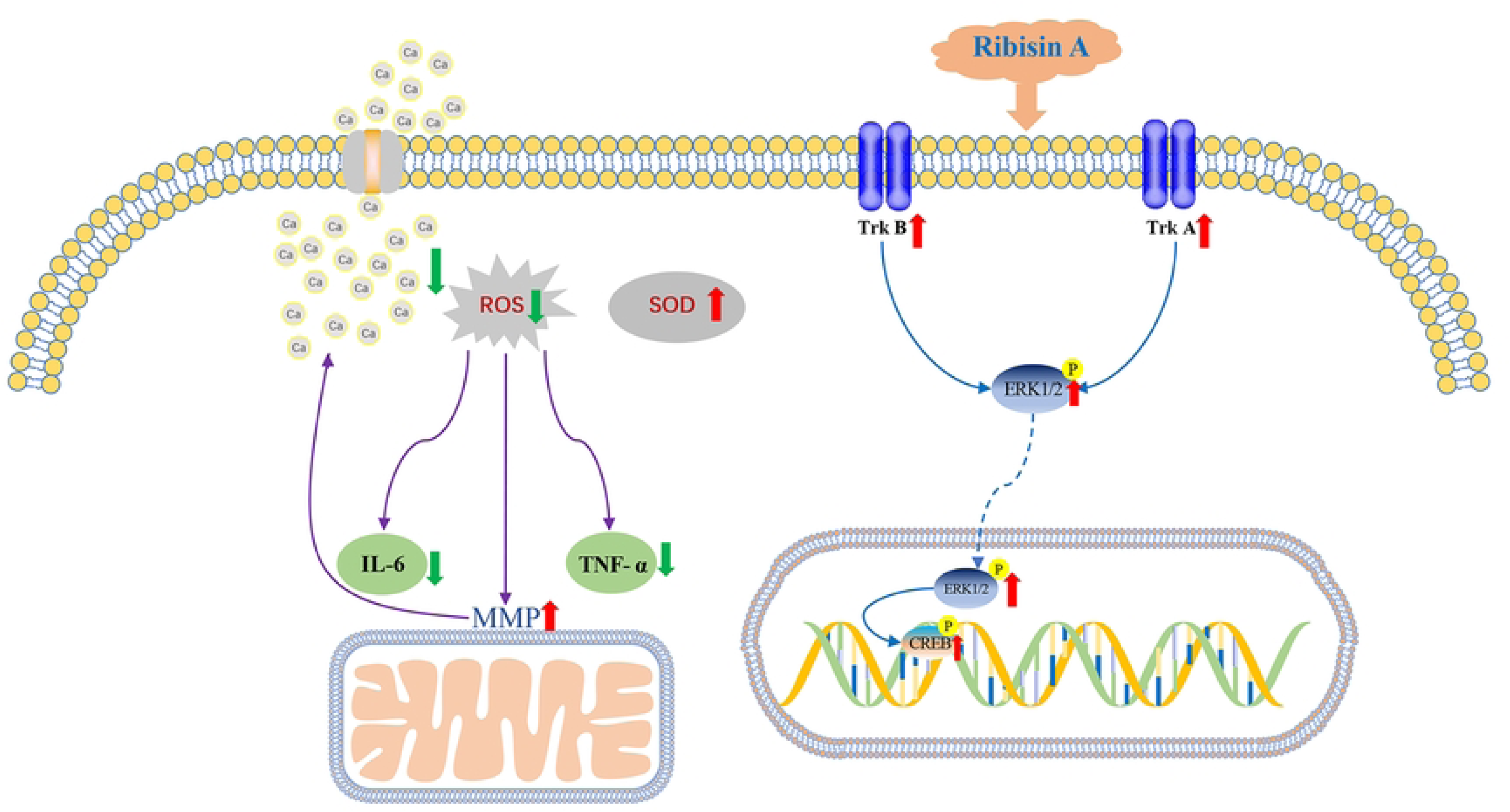
*Neuroprotective mechanism of Ribisin A on H_2_O_2_-induced PC12 cell injury model. Ribisin A could achieve neuroprotective effects by activating the ERK signaling pathway, reducing cellular levels of oxidative stress, inflammatory factors, and apoptosis, and restoring mitochondrial function*.

## Author Contributions

Xin Zhang: Data Curation, Writing of Draft, and Editing. Meng-yu Bao: Data Curation. Jingyi Zhang: Data Curation. Lihao Zhu: Resources. Xin Liu: Formal Analysis. Di Wang: Formal Analysis. Ling-chuan Xu: Supervision. Yu-guo Liu: Supervision. Li-juan Luan: Methodology. Yu-hong Liu: Conceptualization.

## Conflicts of Interest

The authors declare no competing interests.

## Acknowledgements

The authors extend their appreciation to the Shandong University of Traditional Chinese Medicine Experimental Center for their help in Cellular experiments.

## Supplementary information

Supplementary data to this article can be found online at https://doi.org/10.6084/m9.figshare.23559366.

### Abbreviations

ELISA: enzyme-linked immunosorbent assay
MTT: methyl tetrazolium
LDH: lactate dehydrogenase
SOD: superoxide dismutase TNF-α tumor necrosis factor-α
ROS: reactive oxygen species
MMP: mitochondrial membrane potential
AD: Alzheimer’s disease
APP: amyloid precursor protein
BACE-1: beta-site amyloid precursor protein cleaving enzyme-1
AChEI: acetylcholinesterase inhibitors
NMDAR: N-methyl-D aspartate receptor VE Vitamin E
LDH: lactic dehydrogenase
ECL: enhanced chemiluminescence
Ca^2+^: calcium ion
PS: phosphatidylserine
PI: propidium iodide
HO-1: heme oxygenase
NQO1: NAD(P)H quinone reductase 1

## Notes

### Competing Interest Statement

The authors have declared no competing interest.

